# Light-Responsive MicroRNAs in Human Retinal Tissue are Differentially Regulated by Distinct Wavelengths of Light

**DOI:** 10.1101/2023.02.10.528054

**Authors:** Canan Celiker, Kamila Weissová, Kateřina Amruz Černá, Jan Oppelt, Jana Šebestíková, Petra Lišková, Tomáš Bárta

**Author notes:** Corresponding author: Tomáš Bárta, Department of Histology and Embryology, Faculty of Medicine, Masaryk University, Kamenice 3, 625 00 Brno, Czech Republic.

## Abstract

Retinal microRNA (miRNA) molecules play critical roles in a wide range of processes including cell proliferation, cell death, and synaptic plasticity. Recently they have been shown to regulate crucial processes that are associated with perception of light including visual function, light adaptation, and control of genes regulating circadian light entrainment. Despite extensive work on retinal miRNAs in different model organisms, light-regulated miRNAs in human retina are not known. Here, we aim to characterize these miRNAs. We generated light responsive human retinal organoids that express miRNA families and clusters typically found in the retina. Using in-house-developed photostimulation device, we found that 51 miRNAs are up- or downregulated upon brief photostimulation periods. Clustering analysis revealed that only two miRNA families and three clusters are upregulated, while eight families and ten cluster are downregulated upon photostimulation. Additionally, we found that the light-regulated miRNAs have rapid turnover, and their expression is differentially regulated by distinct wavelengths of light. This study demonstrates that only a small subset of miRNAs is light-responsive in human retinal tissue and the generated human retinal organoids are a valuable model for studying the molecular mechanisms of light perception in the retina.

## Introduction

The retina is a specialized, light-sensitive neural tissue that lines the inner surface of the eye. It is composed of several layers of neurons that convert incident light into electrical signals, which are transmitted to the brain for visual perception. The outermost layer of the retina, the photoreceptor layer, comprises two distinct types of photoreceptor cells: rods and cones. Rods are primarily responsible for vision under low-light conditions, while cones are responsible for color vision and visual acuity during daylight conditions. The retina also contains other specialized cell types including bipolar, horizontal, retinal ganglion and Müller cells. The development and function of the retina is intricately regulated by various factors including hormones, growth factors, transcription factors and miRNA molecules. Recent research has demonstrated that miRNAs play a crucial role in regulating light-induced processes within in mice retina, including adaptation to different light intensities and regulation of circadian genes^1,2^, however nothing is known about retina-specific miRNAs in human retina.

miRNAs represent a class of small non-coding regulatory RNAs of 18–24 nucleotides in length found in all metazoans.^3^ They act as post-transcriptional regulators of gene expression by maintaining cellular and tissue homeostasis, controlling development, and signaling pathways.^4^ Expression of miRNAs appears to be highly influenced by the tissue specificity representing the key regulatory elements of virtually all cell processes underlying a cell function. Notably, many miRNAs are specifically expressed in the retina, giving rise to a unique miRNA expression profile in retinal tissue.^3,5,6^

It has been established that miRNAs play critical roles in regulating the expression of genes essential for photoreceptor development and function.^2^ Additionally, they serve as crucial regulators of light-sensitive processes within the retina. In mammals, light regulates the amplitude and duration of the daytime by modulating the activity of several key light-sensitive proteins, including the opsins and melanopsin.^7-9^ Several studies have demonstrated that miRNAs exhibit differential expression in the retina in response to light exposure.^1,3^ One well-studied group of light-regulated miRNAs is the miR-182/183/96 cluster, containing three miRNAs: miR-182, miR-183, and miR-96. This cluster is upregulated in the retina during light adaptation and its overexpression leads to increased cell proliferation and neurogenesis.^1,10^ Additionally to its role in neurogenesis, miR-183 also plays a role in protecting the retina from light-induced damage by reducing oxidative stress through targeting and degrading the key protein SIRT1.^11^ Moreover, the miR-182/183/96 cluster is also implicated in the development of retinal pigment epithelium and circadian rhythms regulation.^3,12,13^

Despite the recognized importance of light-regulated miRNAs in the retina, their functions are not yet fully understood, particularly in humans. This is due to the difficulty of studying miRNAs in the human retina, as the availability of human retinal tissue for experimental procedures is limited. Retinal organoids, which mimic the cellular composition and function of the human retina, represent a promising model for studying light-regulated miRNAs in humans.

Here we employed retinal organoids derived from human pluripotent stem cells as a tool to closely investigate light-regulated miRNAs in the human retina. We developed a device capable of photostimulating retinal organoids using various intensities and wavelengths of light. By utilizing this device, we photostimulated retinal organoids and identified 51 light-regulated miRNAs. Further analysis revealed that these miRNAs exhibited a rapid turnover and differential responses to distinct wavelengths of light. Our findings indicate that human retinal organoids represent a valuable tool for studying light-regulated miRNAs and open new avenues for research on novel light-regulated miRNA targets, the adaptation of the human retina to light, and the regulation of light-regulated circadian rhythms in humans.

## Results

### Generation of light-responsive human retinal organoids

In the first experiment, we aimed to determine the stage of the retinal organoid differentiation process that is applicable for studying light-regulated miRNAs. We generated light-responsive retinal organoids from human induced pluripotent stem (hiPS) cells, as previously described **(Figure 1A)**.^14,15^ We harvested retinal organoids at different stages of the maturation process (D90, D120, D150, and D180) and assessed the expression of opsins (*OPN1SW, OPN1MW, OPN1LW*, and *RHODOPSIN*) and miRNAs (miR-96-5p, miR-182-5p, and miR-183-5p) that have been previously shown to be expressed in the retina.^16-18^ We observed a significant upregulation of opsins at D150 and a significant upregulation of the selected miRNAs from D120 to D180, when compared to D90 **(Figure 1B)**.

**Figure 1:**
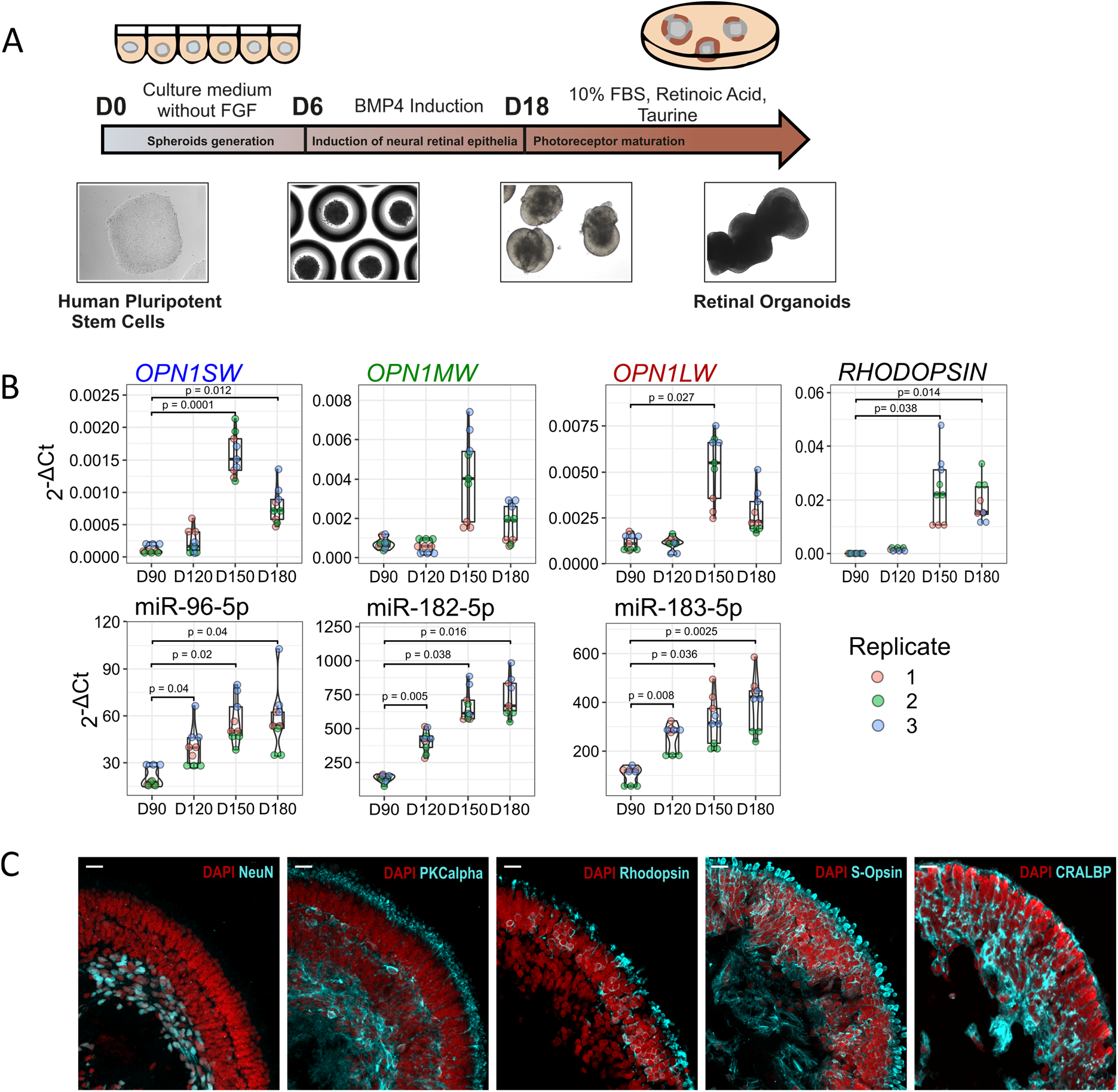
Characterization of retinal organoids. **A)** Schematic presentation of the differentiation protocol. **B)** Expression of OPN1SW, OPN1MW, OPN1LW, RHODOPSIN, and retina-specific miR-96-5p, miR-182-5p, miR-183-5p during retinal organoid differentiation, as determined by RT-qPCR. C) Expression of NeuN, PKCalpha, Rhodopsin, S-Opsin, and CRALBP, as demonstrated using immunofluorescence staining. Scale bars = 20 μm.

Next, we aimed to determine whether the generated retinal organoids contain major cell types typically found in the retina. We performed indirect immunofluorescent staining on cross-sections (at D180). The generated retinal organoids contained a laminated retina, retinal ganglion cells (NeuN+), bipolar cells (PKCα+), photoreceptors (Rhodopsin+, S-Opsin+), and Müller glia (CRALBP+) **(Figure 1C)**. Taken together, our data indicate that the generated retinal organoids between D150-D180 express opsins, retina-specific miRNAs, and contain major cell types typically found in the retina. Therefore, all analysis in this study were done using organoids between D150-D180 of the differentiation process.

### Retinal organoids express retina-specific miRNAs

We conducted next-generation sequencing (NGS) of miRNAs to identify miRNA molecules expressed in retinal organoids. We compared the generated NGS data with a previously published list of retina-specific miRNAs.^19^ We detected 56 miRNAs (out of 60) that have been previously reported to be expressed in the developing retina **(Figure 2A)**. These 56 miRNAs represent approximately 67% of all mapped miRNA reads in human retinal organoids. Since a significant proportion of miRNA genes occur in defined clusters and typically act as families with the same seed sequence, we clustered all detected miRNAs into miRNA clusters and families. This analysis revealed that retinal organoids express miRNA families and clusters that are often involved in regulating neuronal or retinal cell fate **(Figure 2B)**. We identified these miRNA families to be most abundant: mir-9 (involved in neuronal differentiation,^20^), let-7 (promotes terminal cell differentiation,^21,22^), mir-7 (promotes photoreceptor differentiation,^23^), mir-26 (controls neuronal differentiation,^24^), mir-125 (regulates neuronal differentiation and proliferation,^25^). The most abundant miRNA clusters were the miR-7/1179/3529, let-7a/d/f, and miR-183/96/182 clusters (over 15% of total mapped miRNA reads). Members of these clusters have been previously shown to be expressed in the retina or neural tissues.^16-18^ Taken together, these data indicate that retinal organoids possess a miRNA transcriptome that is similar to that of the developing retina.

**Figure 2:**
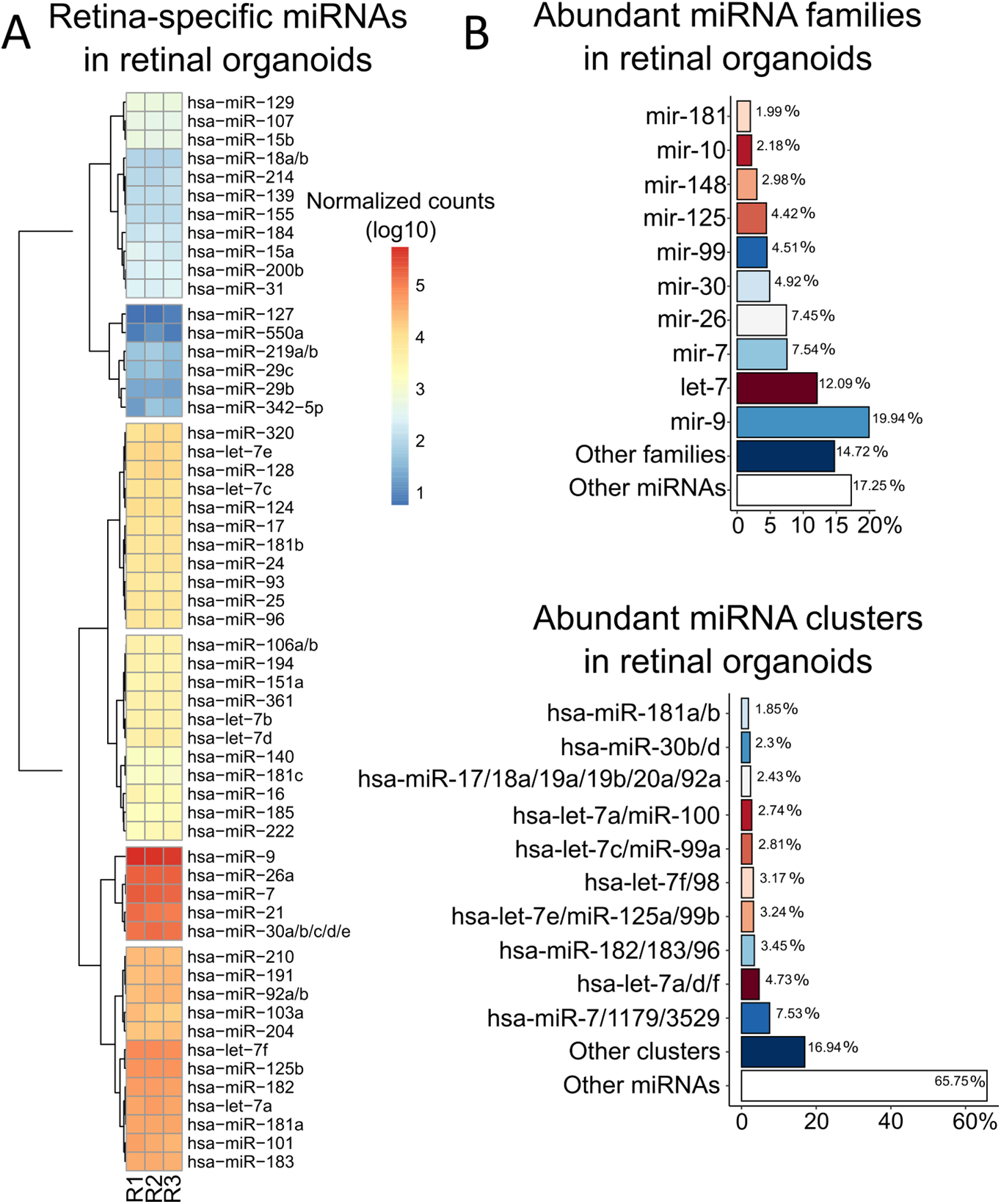
Retinal organoids express retina-specific miRNAs. **A)** Heatmap shows the expression of retina-specific miRNAs in human retinal organoids. R1, R2, and R3 represent different replicates. **B)** Abundance of miRNA families and clusters present in retinal organoids.

### Retinal organoids are responsive to photostimulation at miRNA transcriptional level

To study the effects of photostimulation on miRNA expression in human retinal organoids, we developed Cell LighteR, a photostimulation system capable of controlled, high-throughput, and versatile stimulation of cells cultured in 96-well plates with superior temporal precision. The photostimulation platform consists of a control unit containing a power supply and an Arduino (https://www.arduino.cc/) microcontroller that drives individual light emitting diodes (LEDs) **(Figure S1A, D)**. The control unit is connected to the LED module using a thin wire, and the LED module is placed into an incubator. Emitted light from the LED array in the LED module is directed to individual wells of a black 96-well plate, ensuring that it does not influence neighboring wells **(Figure S1B)**. The plate containing organoids is positioned on top of the LED module and covered with a black lid. The device allows for high-throughput experiments, as it can control up to 3×96-well plates (288 wells) simultaneously. Each LED is individually addressable, allowing for easy, precise, and simultaneous modulation of multiple parameters such as luminous intensity, frequency, and three distinct wavelengths of monochromatic light to deliver adjustable light outputs to organoids in individual wells **(Figure S1C)**. Cell LighteR is equipped with RGB LED chips (type 5050), capable to emit distinct wavelengths of monochromatic light in the red, green, and blue portions of the spectrum [emitting maximum RED (620-625 nm); GREEN (522-525 nm); BLUE (465-467 nm)], allowing for the study of the contribution of individual parts of the spectrum to the effects of photostimulation **(Figure S1C, S1E)**.

To ensure that photostimulation of retinal organoids is performed at a physiological level and does not induce apoptosis or overheating of the organoid culture, we conducted a series of experiments to determine the appropriate level of luminous intensity. These experiments included assessing the expression levels of genes involved in apoptosis (*PUMA, BIM*) upon photostimulation and measuring changes in the temperature of the culture media during the photostimulation process. Organoids were placed in a black 96-well plate containing cell culture media and positioned on a LED module inside an incubator. The organoids were photostimulated using different luminous intensities (15, 50, 500, and 4200 lx) of white light and harvested 3 hours after the start of the photostimulation process. We did not observe a significant upregulation of *PUMA* expression at 500 and 4200 lx, and *BIM* expression remained unchanged across all luminous intensities tested **(Figure S1G)**. Therefore, for short-term photostimulation of retinal organoids, we used a luminous intensity of 100 lx. To determine changes in temperature during photostimulation, we utilized a digital thermometer equipped with ultra-thin sensors (Dallas Instruments) to continuously monitor temperature. One sensor was immersed in the culture media in a 96-well plate that was being photostimulated, while the second sensor was placed in another 96-well plate kept in the dark. The temperature difference did not exceed 0.3°C during the measurement, indicating that there was no overheating of the cell culture media during the photostimulation process **(Figure S1F)**.

Having determined the miRNA expression profile of human retinal organoids and the physiological level of luminous intensity for the photostimulation process, we aimed to identify miRNA molecules that are light-responsive. We photostimulated retinal organoids for 1 hour and 3 hours respectively, using white light at 100 lx of luminous intensity and performed miRNA NGS. Principal component analysis of the NGS data revealed that the miRNA expression profile differs between the dark and light conditions, while there is almost no difference between 1 hour and 3 hours time-points of photostimulation, and the replicates clustered together **(Figure 3A)**. Differential expression analysis revealed 51 miRNAs that were significantly up- or down-regulated (p_adj_ < 0.05 in both time-points) upon photostimulation **(Figure 3B)**, while there was no induction of apoptosis, as demonstrated by non-significant change in the expression of genes involved in apoptosis or DNA damage (*BIM, BAX, BCL2, PUMA*, and *TP53*) **(Figure S2)**. Clustering analysis revealed that only two miRNA families (mir-124 and mir-216) and three miRNA clusters (miR-182/183/96, miR-216a/217, and miR-29a/29b-1/29b) were significantly upregulated **(Figure 3C)**, while eight miRNA families and ten clusters were downregulated upon photostimulation **(Figure 3D)**. Functional annotation revealed that the significantly upregulated miRNAs play roles in multiple cellular functions including neural differentiation, cell proliferation, DNA damage, and circadian rhythms **(Figure 3E)**.

**Figure 3:**
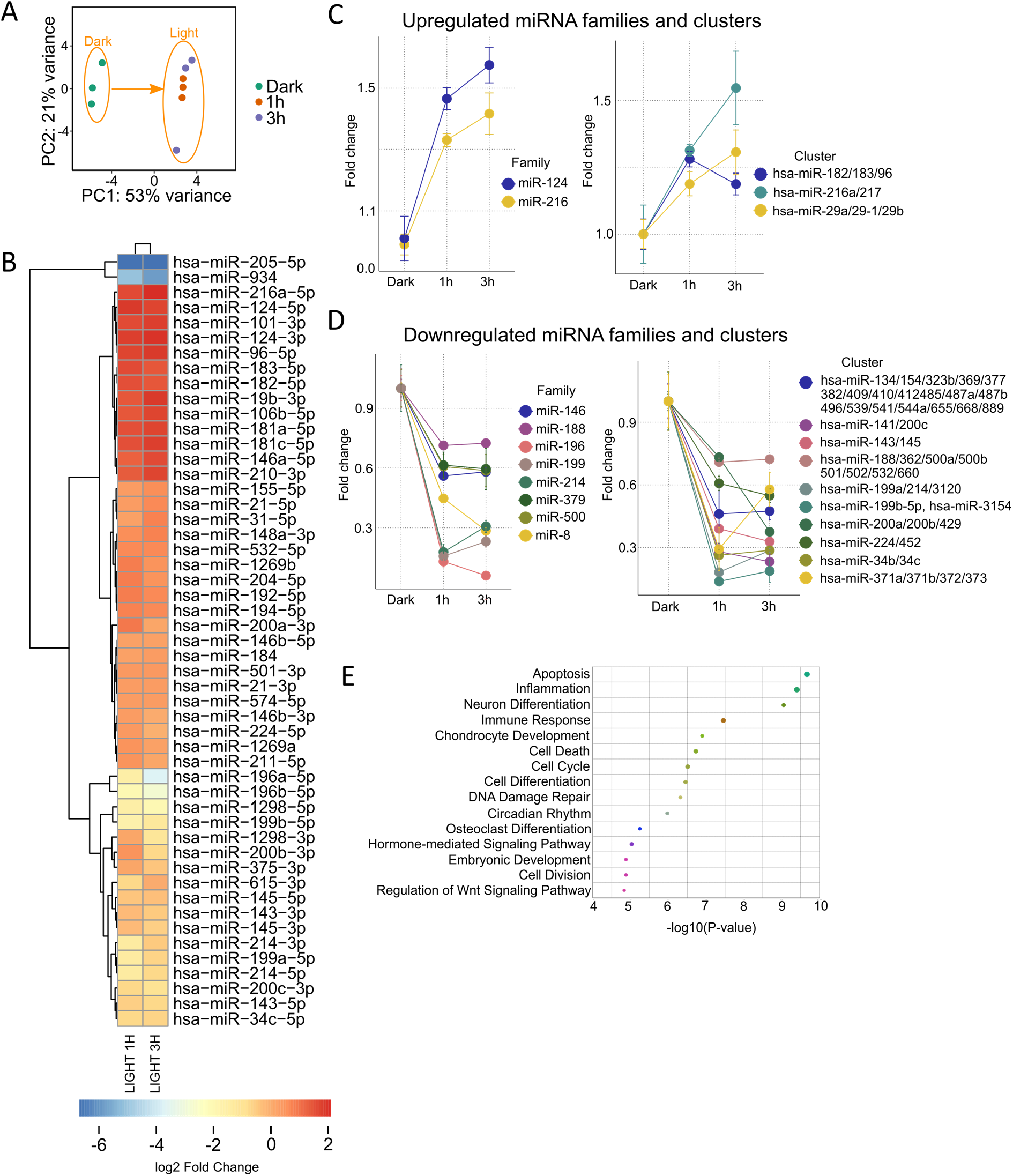
Retinal organoids demonstrate transcriptional responsiveness to photostimulation at the miRNA level. **A)** Principal component analysis (PCA) of miRNA expression profiles in retinal organoids following 1 hour and 3 hours of photostimulation. **B)** Heatmap illustrates miRNAs that were significantly up- or down-regulated upon photostimulation. **C)** Analysis of miRNA families and clusters that are significantly down-regulated upon photostimulation. **D)** Analysis of miRNA families and clusters that are significantly up-regulated upon photostimulation. **E)** Functional annotation of all significantly up-regulated miRNAs upon photostimulation, as determined using the TAM 2.0 tool.

Taken together, only a small number of miRNA molecules are differentially expressed upon photostimulation in retinal organoids, suggesting that only a small subset of miRNA clusters and families are involved in the reaction of the human retina to light.

### Light-regulated miRNAs have rapid turnover and respond to different wavelengths of light

Since a small subset of miRNAs are responsive to light, we aimed to select representative miRNA molecules for detailed expression analysis. We selected nine different miRNAs that were shown to be up- or down-regulated upon photostimulation: miR-96-5p, miR-182-5p, and miR-183-5p representing the retina-specific miR-182/183/96 cluster; miR-204-5p and miR-211-5p as members of the mir-204 family; miR-145-5p as a member of the miR-143/145 cluster; miR-196a-5p as a representative of the mir-196 family; miR-214-3p as a member of the mir-214 family; and miR-205-5p as the most down-regulated miRNA upon photostimulation.

We aimed to assess the expression turnover of the selected miRNAs during alternating periods of darkness and photostimulation. Retinal organoids were cultured in darkness for 24 hours and then exposed to white light at a luminous intensity of 100 lx for 60 minutes. The light was then switched off and the organoids were cultured in darkness for an additional 60 minutes. Samples were harvested at 15-minute or 30-minute intervals, as depicted in **Figure 4A**. The miR-182/183/96 cluster, miR-204-5p, and miR-211-5p were gradually upregulated during the photostimulation period, with the highest expression observed at the 60-minute time point. Upon switching off the light, expression gradually decreased, returning to levels observed in darkness after 30 minutes (**Figure 4A**). Conversely, miRNAs that were shown to be downregulated upon photostimulation, including miR-205-5p, miR-214-3p, miR-196a-5p, and miR-145-5p, were gradually downregulated and reached their lowest expression between 30-60 minutes of photostimulation. Upon switching off the light, expression of these miRNAs returned to their original levels as observed in darkness (**Figure 4A**). These results indicate that the light-regulated miRNAs have rapid turnover in the human retina.

**Figure 4:**
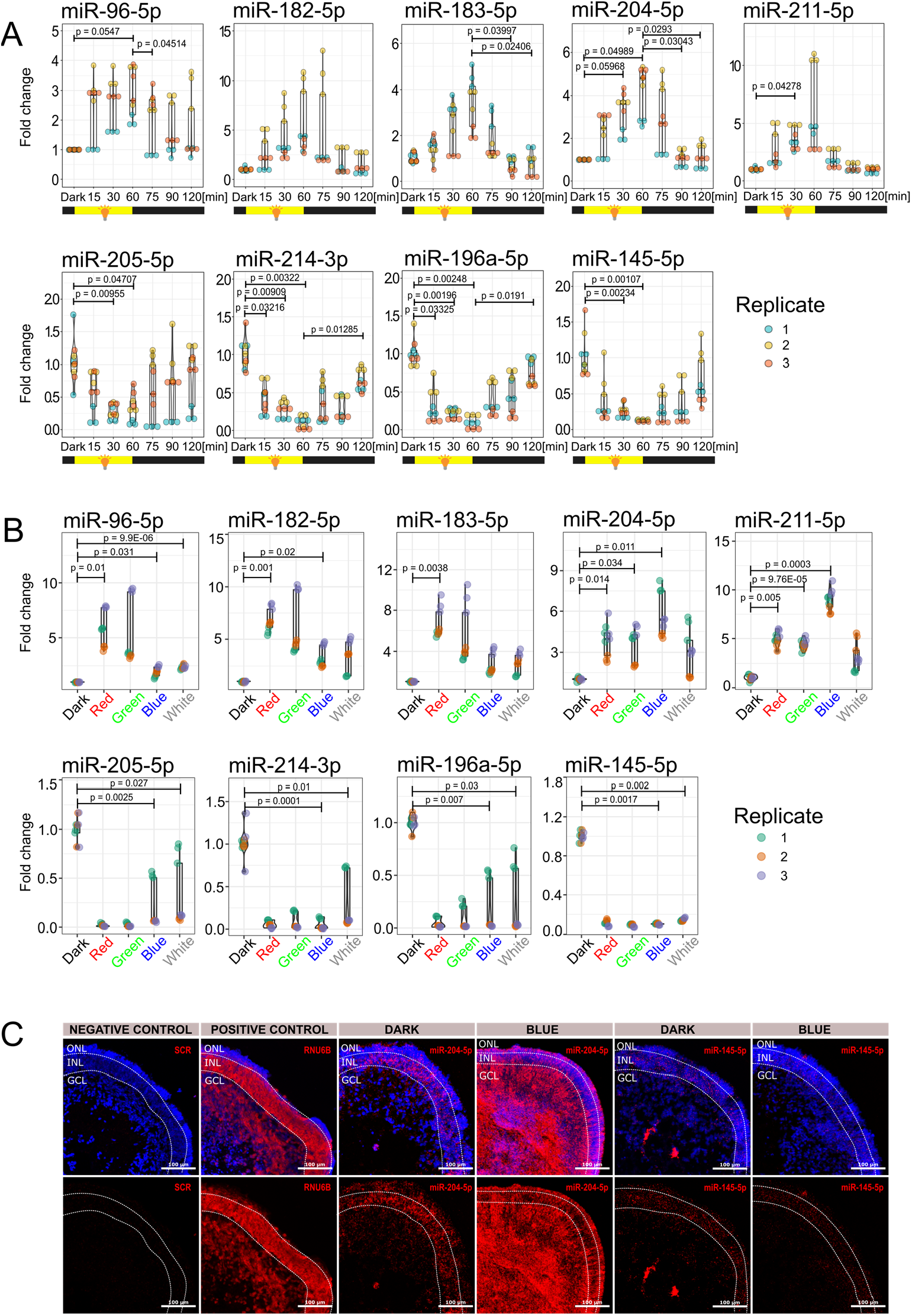
Light-regulated miRNAs display rapid turnover and exhibit distinct responses to different colors of light. **A)** Expression of miRNAs during alternating periods of darkness and light, as determined by RT-qPCR. **B)** Expression of miRNAs upon photostimulation using different wavelengths of light at 100 lx for 1 hour, as determined by RT-qPCR. **C)** miRNA *in situ* hybridization upon exposure to blue light at 100 lx for 1 hour, as determined by miRNA scope. ONL-presumptive Outer Nuclear Layer, INL – presumptive Inner Nuclear Layer, GCL – presumptive Ganglion Cell Layer.

There is an intriguing question as to whether these miRNAs respond differently to distinct wavelengths of light. To address this question, we photostimulated retinal organoids using distinct wavelengths of light [RED (620-625 nm), GREEN (522-525 nm), and BLUE (465-467 nm)] **(Figure S1B)**. As each channel in the RGB LED has a different light power **(Figure S1C)**, we adjusted individual channels to a luminous intensity of 100 lx. Interestingly, red and green light were identified as the strongest inducers of the members of the miR-182/183/96 cluster, while blue light was the strongest inducer of the mir-204 family, as both members, miR-204-5p and miR-211-5p, were upregulated using blue light. Surprisingly, we did not observe any strong association between the level of downregulation of miRNAs and the color of light, as all miRNAs (miR-205-5p, miR-214-3p, miR-196a-5p, miR-145-5p) were downregulated in all conditions.

Next, we aimed to evaluate the spatial expression of miRNAs in retinal organoids using *in situ* miRNA hybridization (miRNA scope). Intriguingly, blue light had the greatest effect on the upregulation of the mir-204 family members and on the downregulation of the miR-145-5p (**Figure 4B**), therefore we aimed to determine the localization and expression of the miR-204-5p and the miR-145-5p in retinal organoids photostimulated using blue light (100 lx for 1 hour). In the dark condition, both miRNAs were predominantly localized in the presumptive Inner Nuclear Layer (INL). Upon photostimulation, miR-204-5p expression was highly upregulated in the presumptive Ganglion Cell Layer (GCL) and Outer Nuclear Layer (ONL), while miR-145-5p expression was downregulated in the INL (**Figure 4C**). These findings suggest that retinal ganglion cells and photoreceptors upregulate miR-204-5p expression, while miR-145-5p expression decreases in cells present in the INL.

## Discussion

In this study, we identified miRNAs that respond to photostimulation in the human retina using retinal organoids as a model. We showed that retinal organoids express a number of retina-specific miRNA families and clusters that have been previously associated with the regulation of neuronal or retinal cell fate, including miR-9, let-7, miR-7, miR-26, and miR-125.^19^ To study the light-regulated miRNAs, we developed a photostimulation system Cell LighteR and used it to stimulate the organoids with light of different wavelengths. We showed that retinal organoids are responsive to photostimulation at the miRNA transcriptional level, as evidenced by the altered expression of multiple miRNA families and clusters in response to light exposure. Furthermore, our research revealed that many miRNAs in the retina have a high turnover rate, much faster than miRNAs expressed in other cell types or tissues.^26^ Importantly, some miRNAs exhibit distinct responses to various wavelength ranges of light. In particular, exposure to red and green light strongly induced expression of the miR-183/182/96 cluster, while blue light exposure upregulated the expression of the members of the miR-204 family.

To the best of our knowledge, our study is the first to report light-regulated miRNAs in human retinal organoids, providing new insights into the regulation of light-induced transcription and light adaptation in the human retina. However, some light-responsive neuronal miRNAs have already been identified in mice.^26,27^ Krol et al^26^ identified neuronal miRNAs that are downregulated during dark adaptation and upregulated in light-adapted mice retinas. Surprisingly, despite using multiple approaches, including Exiqon arrays and Illumina sequencing, they identified only five light-responsive miRNAs (miR-96, miR-182, miR-183, miR-204, and miR-211 - both members of the mir-204 family). However, here we identified over 10-fold more light-responsive miRNAs. The striking difference can be explained by **I)** the deeper sequencing used in our study with the number of miRNA reads per sample ranging from 1.7 to 3.1 million (**Fig. S3)**; **II)** different experimental setups; **III)** diverse experimental models. Despite the difference between the number of miRNAs identified, the miR-183/182/96 cluster and the mir-204 family were recognized as light-regulated miRNAs in both studies. Here we provide much comprehensive list of light-regulated miRNAs in humans that includes a detailed clustering, functional annotation analysis, expression turnover assessment, and analysis of the effects of different wavelengths of light on light-regulated miRNAs, further expanding our knowledge about light-regulated miRNAs in human retina.

Previously, our lab demonstrated that the miR-182/183/96 cluster is an important morphogenetic factor in human retinal organoids that targets *PAX6* expression.^10^ Other studies have shown that this miRNA cluster is involved in the development of photoreceptor outer segments, synapses and retinal laminar structure.^28-31^ Furthermore, the voltage-dependent glutamate transporter Slc1a1 is targeted by the miR-182/183/96 cluster, and under low glutamate conditions, Slc1a1 can clear glutamate from the photoreceptor synaptic area. This miRNA cluster is also implicated in maintaining cone outer segments by regulating genes related to membrane trafficking, lipid metabolism, and cilia formation.^32^ Given the critical role of the miR-182/183/96 cluster in the retina and its responsiveness to light, it is possible that light stimulation will play an important role in shaping the post-natal retina.

The miRNA oscillatory pattern is a highly conserved mechanism and has significant consequences for circadian timing.^33^ Here we identified nine light-regulated miRNAs - hsa-mir-96, hsa-mir-194-2, hsa-mir-181a-1, hsa-mir-194-1, hsa-mir-182, hsa-mir-183, hsa-mir-192, hsa-mir-181a-2, and hsa-miR-124 - that are linked to the regulation of circadian rhythms.^3,34^ Besides well-known cluster miR-182/183/96 that targets critical circadian genes including *ADCY6, CLOCK, BMAL1*,^3,35^ the miR-194/192 cluster has been found to target other circadian genes including *PER1, PER2*, and *PER3*.^36,37^ Interestingly, the miR-124 represents an important regulator of *Drosophila* circadian locomotor rhythms and under the dark conditions flies lacking miR-124 (miR-124^*KO*^) have a greatly enhanced circadian behavior phase.^38^ Similarly, it has been shown that miR-124 mutants’ advanced circadian phase functionally connects with the suppression of miR-124 targets in BMP signaling.^39^ Although some light-regulated miRNAs are regulated independently of the circadian rhythms,^26^ they may still play a critical role as circadian regulators that can quickly respond to rapidly changing light stimuli.

The quick turnover of miRNAs in retinal neurons and the resulting fast changes in target gene expression can enhance the precise encoding of visual stimuli in a rapidly changing environment.^31^ Our study provides new insights into the regulation of miRNA expression in response to light in the human retina. The discovery of wavelength-specific miRNA responses and the high turnover rate of miRNAs in the retina highlights the importance of deeper sequencing methods in identifying miRNAs, and can guide future research on the functional analysis of these miRNAs to understand their roles in the regulation of light-induced transcription and light adaptation in the human retina. Light regulated miRNAs have significant implications for the formation and function of the retina, as well as for circadian timing. This work paves the way to study the complex interplay between light and miRNA regulation in the retina and its effects on visual function and circadian rhythms.

## Material and Methods

### Data and code availability

Bulk miRNA sequencing data have been deposited at GEO and are currently publicly available. Accession number is: GSE223379.

This paper does not report original code.

Any additional information required to reanalyze the data deposited in this paper is available from the lead contact upon request.

### hiPS cell lines

Neonatal dermal fibroblasts (purchased form Lonza, Product code: CC-2509) were reprogrammed using an Epi5™ Episomal iPSC Reprogramming Kit (Invitrogen) according to the manufacturer’s instructions. hiPS cells (Neo5) were maintained in Essential 8 medium (Gibco) on Vitronectin-coated dishes and regularly passaged every ∼4-5 days. hiPS cells between passages 40-60 were used in this study. Characterization is the cell line was done using flowcytometry analysis of SOX2 and OCT3/4 expression (**Figure S4**).

### Retinal organoids generation

Retinal organoids were generated using the previously reported protocols ^14,15^ with slight modifications. HiPS cells cultured in Essential 8 medium (Gibco) were seeded into a U-shaped, cell-repellent 96-well plate (5000 cells/well), (Cellstar). After 48 hours (day 0 of the differentiation process), the culture medium was changed to growth factor-free Chemically Defined Medium (gfCDM) containing 45% Iscove’s modified Dulbecco’s medium (IMDM, Gibco), 45% Ham’s F12 (F12, Gibco), 10% KnockOut Serum Replacement (Gibco), 1% chemically defined lipid concentrate (Gibco), 1% Penicillin-Streptomycin Solution (Biosera), 10 μM β mercaptoethanol (Sigma-Aldrich). On day 6, recombinant human BMP4 (Peprotech) was added to the culture to the final concentration 1.5 nM and then the medium was changed every third day. On day 18 of the differentiation process, gfCDM was changed to a NR medium containing DMEM/F12 (Gibco), 1% N-2 supplement (Gibco), 1% GlutaMAX supplement (Gibco), 10% foetal bovine serum (FBS; Biosera), 0.5 mM retinoic acid (Sigma), 0.1 mM Taurine (Sigma), 1% Penicillin-Streptomycin Solution (Biosera). The organoids were cultured in 96-well plates to day 18 and then they were transferred to 10 cm Petri dish.

### Photo-stimulation of retinal organoids

Twelve hours before photo-stimulation retinal organoids were transferred to black 96-well plate with clear round bottom, ultra-low attachment (Corning) into the fresh culture media (one organoid per well). The LED module was placed into a conventional tissue incubator and the 96-well plate containing organoids was positioned onto the LED module and covered with the black lid. Source code that controls individual LEDs was compiled and uploaded into Arduino UNO microcontroller (https://www.arduino.cc) (**Figure S1**).

### RNA isolation

At least 6 organoids per sample were washed with phosphate-buffered saline (PBS), and were homogenized using a 1 ml insulin syringe in 300 μl RNA Blue Reagent (an analogue of Trizol) (Top-Bio). After 1-bromo-3-chloropropane (Sigma-Aldrich) addition and centrifugation (21000 g, 15 minutes, 4 °C), the cell lysates were separated into three layers. The top aqueous layer containing RNA was transferred to a new tube and precipitated by addition of isopropanol, followed by 10 minutes incubation on ice and centrifugation (18000 g, 10 minutes, 4 °C). RNA pellets were washed with 70% ethanol (8500 g, 10 minutes, 4 °C), air-dried, solubilized in PCR grade water (Top-Bio). For RT-qPCR, after organoids were homogenized using a 1 ml insulin syringe in 300 μl RNA Blue Reagent, Direct-zol™ RNA Microprep kit (Zymo Reseach) was used according to the manufacturer’s instructions and as described previously.^40^

### RT-qPCR analysis

RNA was reverse transcribed using a High-Capacity cDNA Reverse Transcription Kit (Applied Biosystems). The RT product was amplified by LightCycler 480 Real-Time PCR system (Roche) using PowerUp SYBR Green Master Mix (Applied Biosystems). Primer sequences are shown in Supplementary Table S1. Data sets were normalized to the corresponding levels of GAPDH mRNA. For the determination of miRNA expression, RNA was reverse transcribed (16°C, 30 minutes; 42°C, 30 minutes; 85°C, 5 minutes) using a TaqMan MicroRNA Reverse Transcription Kit (Applied Biosystems) using specific primers (Applied Biosystems), (Supplementary Table S2). Reverse transcription products were then amplified by LightCycler 480 Real-Time PCR system (95°C, 5 minutes; 95°C, 15 seconds; 60°C, 60 seconds; 40 cycles) using TaqMan Universal PCR Master Mix and specific probes for miRNAs (Applied Biosystems), (Supplementary Table S2). Relative microRNA expression was determined using the ΔΔCt method and normalized to endogenous control *RNU6B*.

### Small RNA library preparation and sequencing

RNA quality was assessed by TapeStation 2200 (RNA Screen Tape; Agilent Technologies), and only samples with RINe values ≥ 9 were used for library preparation. NEBNext Multiplex Small RNA Library Prep Set for Illumina (New England Biolabs) was used to prepare libraries for further sequencing according to the manufacturer’s instructions. Briefly, 800 ng of total RNA were used to create size selected small RNA libraries (size selection with 6% PAGE gel). The sequencing was performed with 2.0 pM library using the NextSeq® 500/550 High Output Kit v2.5 (75 cycles; Illumina).

### miRNAScope assay and immunohistochemistry

The organoids were fixed in 4% paraformaldehyde for 30 minutes at room temperature, washed three times with PBS, cryopreserved in 30% sucrose (Sigma Aldrich) in PBS for overnight at 4°C, embedded in Tissue-Tek O.C.T. Compound medium (Sakura), and sectioned on a cryostat (Leica CM1850) (∼10 μm) using Superfrost Plus™ Adhesion Microscope slides (Epredia).

In situ hybridization (ISH) assay was performed using a miRNAscope™ HD (Red) Assay (ACDBio) following the fixed-frozen tissue sample protocol provided by the manufacturer. Briefly, the slides were washed with PBS to remove O.C.T. and treated with hydrogen peroxide (ACDBio) for 10 min at room temperature, followed by 10-fold diluted (in PBS) Protease III treatment for 40 min at 40°C in a HybEZ Oven (ACDBio). Hybridization was performed using probes from ACDBio: SR-RNU6-S1 (positive control), SR-Scramble-S1 (negative control), SR-mmu-miR-204-5p-S1 (target) and SR-hsa-145-5p-S1 (target). Signal amplification and detection reagents (ACDBio) were applied sequentially and incubated in AMP 1, AMP 2, AMP 3, AMP 4, AMP 5, and AMP 6 reagents, for 30, 15, 30, 15, 30, 15 min, respectively. Before adding each AMP reagent, samples were washed twice with the washing buffer (ACDBio). After probe hybridization and signal amplification, hybridization signals were visualized by chromogenic reactions using FastRed (ACDBio).

For immunofluorescent staining the sections were washed with PBS and blocked with 0.3% Triton X-100 (Sigma Aldrich), 5% normal goat serum (Sigma Aldrich) in PBS for 1 hour at room temperature in a humidified chamber. The primary antibodies were diluted in Antibody diluent containing PBS with 0.3% Triton X-100 and 1% bovine serum albumin (Sigma Aldrich) and applied to the sections overnight at 4°C in a humidified chamber. The following primary antibodies were used: Rhodopsin (SantaCruz, 1:200), S-Opsin (Abcam, 1:200), NeuN (CST, 1:200), PKC-alpha (SantaCruz, 1:200), and CRALBP (SantaCruz, 1:200). The sections were washed with Antibody diluent and secondary antibodies Goat anti-Mouse IgG Alexa 488 (Invitrogen, 1:1000) and Goat anti-Rabbit IgG Alexa 594 (Invitrogen, 1:1000) were applied for 1 hour at room temperature in a humidified chamber. The nuclei were stained with 1 μg/mL DAPI in PBS for 4 minutes at room temperature and the sections were mounted using Fluoromount Aqueous Mounting Medium (Sigma Aldrich).

### Flow cytometry analysis

hiPS cells were characterized using human pluripotent stem cell transcription factor kit (BD Biosciences). hiPS cells were dissociated using 0.5 mM EDTA in PBS and counted with haemocytometer. Cells were then fixed using cytofix fixation buffer (BD Biosciences) and permeabilized in Perm/wash buffer (BD Biosciences), and stained with Alexa Flour 647 Sox2 and PerCP-Cy5.5 Oct3/4 antibodies and their isotype controls (BD Biosciences) at RT for 30 min. Cells were washed with Perm/wash buffer and resuspended in PBS. Cells were analysed immediately after labelling using BD FACSAria equipped with BD FACSDiva™ software (v. 6.1.3).

### RT-qPCR statistical analysis

All experiments were performed at least three independent replicates, and each replicate was performed in triplicates. Individual data points are shown in each graph. p-values were calculated by the two-tailed Student’s t-test. The corresponding significance level (p-value) are provided in each panel. All comparisons with p-value <0.05 were considered significant.

### Small RNA-seq data processing

The quality of the raw sequencing data was assessed using FastQC (v0.11.9).^41^ Minion and Swan (Kraken package, v16.098)^42^ were used to scan and identify adaptor sequences which were subsequently removed by Cutadapt (v2.5).^43^ Only adapter-containing reads were kept for further processing. The adapter-trimmed reads were further processed using the following steps: 1) removal of very low-quality read ends (Phred<5); 2) keeping only reads with a Phred score of 10 over at least 85% of the length; 3) only reads within 16-27 bp were kept as potential microRNA reads. FASTX-Toolkit (v0.0.14)^44^ was used for the quality filtering; the rest of the steps were performed by Cutadapt (v2.5)^43^ and bash scripting. The quality of the final pre-processed reads was assessed with overall mapping rates to the human reference genome (hg38),^45^ and the general quality of the pre-processed reads was assessed by Bowtie (v1.3.1).^46^ Reads mapping to rRNAs, tRNAs, snoRNAs, snRNAs, or YRNAs^45^ with fewer mismatches than to miRNAs (miRBase; v22.1)^47^ were excluded. The raw microRNA expression levels were quantified by seqcluster (v1.2.8)^48^ and seqbuster/miraligner (v3.5).^49^ R (v3.6.3)^50^ was used for further evaluation and visualization of the data. Differential expression was calculated with DESeq2 (v1.24.0).^51^ All data analysis and visualizations were conducted using R 3.4.458 with installed Bioconductor project.^52^ The following R packages were used to analyze and visualize data: ggplot2,^53^ pheatmap,^54^ RColorBrewer,^55^ ggstatsplot,^56^ and mirbase.db^57^ package was used to cluster and visualize miRNAs into families and clusters.

## Acknowledgement

This publication was supported by the Czech Science Foundation (GA21-08182S), and by Ministry of Health of the Czech Republic, grant nr. NU22-07-00380. PL was supported by national funds from the Ministry of Education, Youth and Sports of the Czech Republic under the European Joint Program for Rare Diseases (Solve-RET 8F20004 No. 825575).

