## Supplementary files for "Light-Responsive MicroRNAs in Human Retinal Tissue are Differentially Regulated by Distinct Wavelengths of Light"

### Supplementary Information

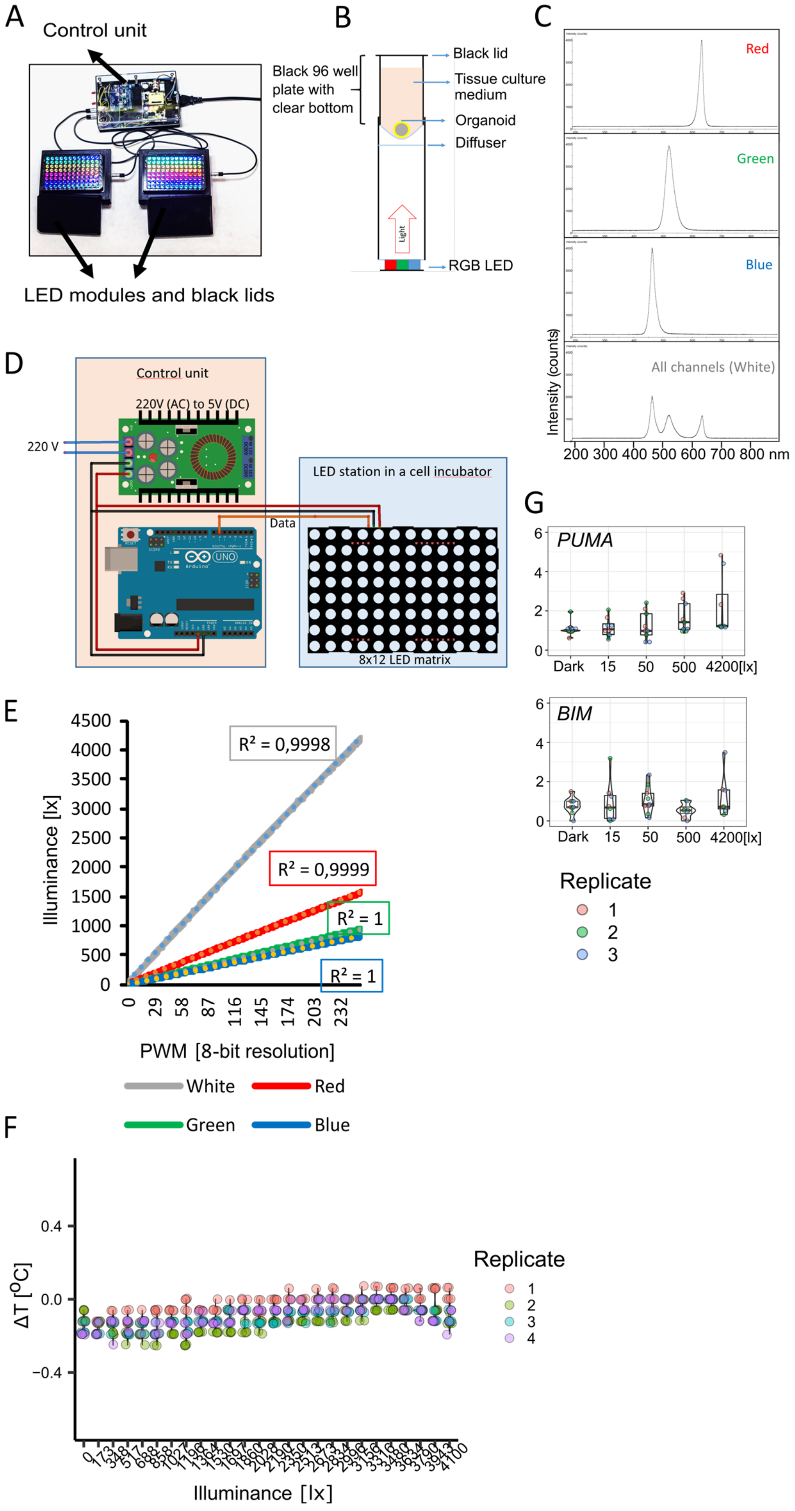


**Supplementary Figure 1: Description of Cell LighteR – a device for photostimulation of human retinal organoids. A)** The Cell LighteR with LED modules. **B)** Schematic illustration of cross-section of a single well in the 96-well plate LED module. **C)** Emission wavelengths of individual LED channels. **D)** Schematic illustration of Cell LighteR. **E)** Luminous intensities of individual LED channels. **F)** Temperature change (ΔT) in a well during photostimulation using different light intensities relative to dark well, as determined by ultra-thin digital sensor (Dallas Instrument). **G)** Expression of *BIM* and *PUMA* genes upon 3 hours of photostimulation using different light intensities, as determined by RT-qPCR.

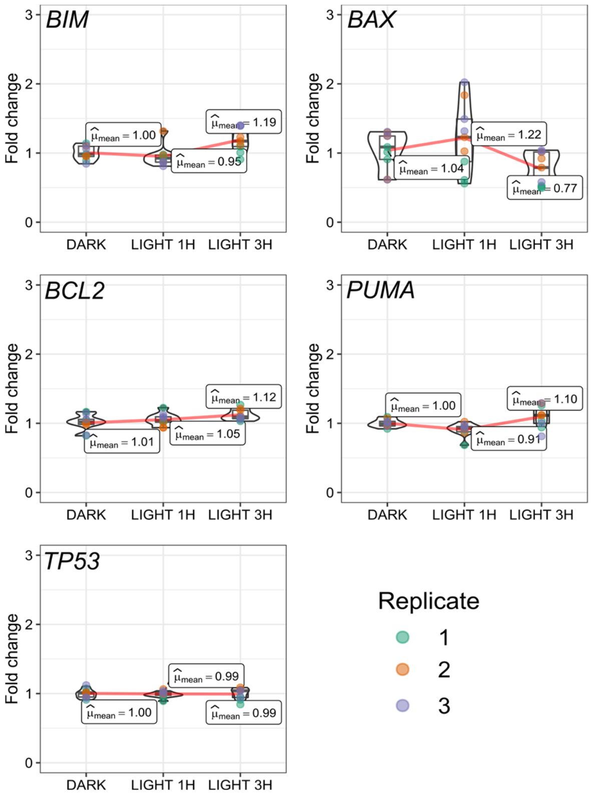


**Supplementary Figure 2: Light stimulation does not upregulate genes involved in apoptosis and DNA damage.** Expression of *BIM*, *BAX*, *BCL2*, *PUMA*, and *TP53* upon 1h and 3h of photo stimulation, as determined by RT-qPCR.

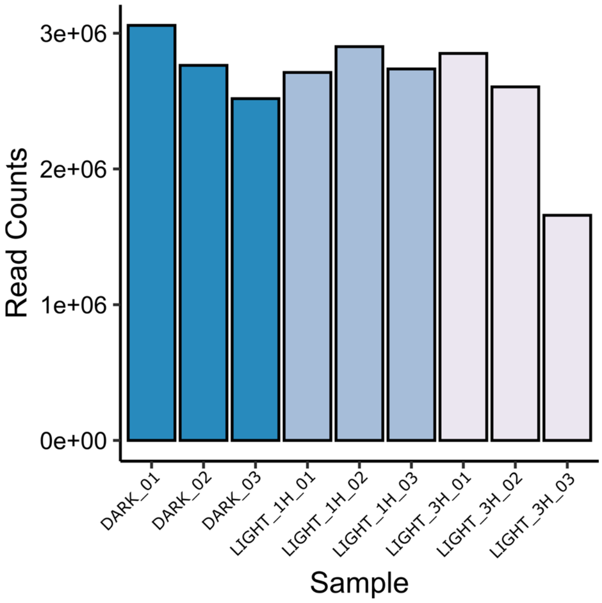


**Supplementary Figure 3: Number of miRNA reads in each NGS sample.**

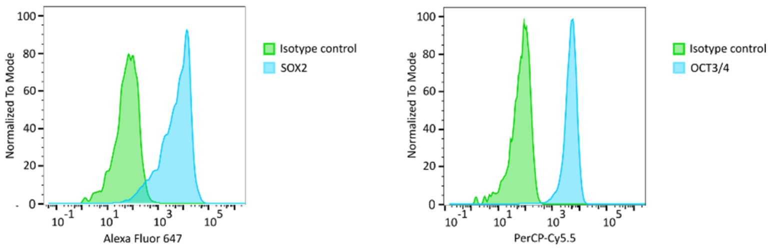


**Supplementary Figure 4:** Characterization of hiPS cell line Neo5 using flowcytometry analysis of SOX2 and OCT3/4 expression.

**Table S1.** List of the primer sequences used for SYBR-green RT-PCR

| Gene Name | Gene Sequence (5'-3') |
|---|---|
| OPN1SW forward | ATACCGCAGCGAGTCCTATAC |
| OPN1SW reverse | GATCCTACCATCACAACCAC |
| OPN1MW forward | CATCTTTGGTTGGAGCAGGTACT |
| OPN1MW reverse | TCTCTGCCTTCTGGGTGGAT |
| OPN1LW forward | GCCTACTTTGCCAAAAGTGC |
| OPN1LW reverse | GATGAGACCTCCGTTTTGGA |
| RHODOPSIN forward | TTTGGAGGGCTTCTTTGCCA |
| RHODOPSIN reverse | CCTCGGGGATGTACCTGGAC |
| PUMA forward | ACGACCTCAACGCACAGTACGA |
| PUMA reverse | CCTAATTGGGCTCCATCTCGGG |
| BIM forward | TTCTGAGTGTGACCGAGAAGG |
| BIM reverse | TGCCTTCAGGATTACCTTGTG |
| BAX forward | TGATGGACGGGTCCGGG |
| BAX reverse | GCAATCATCCTCTGCAGCTC |
| BCL2 forward | CCCGCGACTCCTGATTCATT |
| BCL2 reverse | AGTCTACTTCCTCTGTGATGTTGT |
| TP53 forward | TTCACCCTTCAGATCCGTGG |
| TP53 reverse | AGTCTGAGTCAGGCCCTTCT |
| GAPDH forward | TGCACCACCAACTGCTTAGC |
| GAPDH reverse | GGCATGGACTGTGGTCATGAG |

**Table S2.** List of the miRNA primers/probes used for TaqMan® MicroRNA Assay

| Assay Name | Assay ID | Catalog Number # |
|---|---|---|
| RNU6B | 001093 | 4440887 |
| mmu-miR-96 | 000186 | 4427975 |
| hsa-miR-182 | 002334 | 4427975 |
| hsa-miR-183 | 002269 | 4427975 |
| hsa-miR-204 | 000508 | 4427975 |

| | | |
|---|---|---|
| hsa-miR-211 | 000514 | 4427975 |
| hsa-miR-196a | 241070_mat | 4427975 |
| hsa-miR-205 | 000509 | 4427975 |
| hsa-miR-145 | 000467 | 4427975 |
| hsa-miR-214 | 002306 | 4427975 |
